# *Ex vivo* to *in vivo* model of malignant peripheral nerve sheath tumors for precision oncology

**DOI:** 10.1101/2022.04.29.490078

**Authors:** Himanshi Bhatia, Alex T. Larsson, Ana Calizo, Kai Pollard, Xiaochun Zhang, Eric Conniff, Justin F. Tibbitts, Sara H. Osum, Kyle B. Williams, Ali L. Crampton, Tyler Jubenville, Daniel Schefer, Kuangying Yang, Yang Lyu, Jessica Bade, James C. Pino, Sara J.C. Gosline, Christine A. Pratilas, David A. Largaespada, David K. Wood, Angela C. Hirbe

## Abstract

Malignant peripheral nerve sheath tumors (MPNST) are aggressive soft tissue sarcomas that often develop in patients with neurofibromatosis type 1 (NF1-MPNST), but can occur sporadically. Through a multi-institution collaboration, we have developed 13 NF1-associated MPNST patient-derived xenografts (PDX). Genomic analysis of the PDX-tumor pairs identified somatic mutations in *NF1* (61%), *SUZ12* (61%), *EED* (15%), and *TP53* (15%), and chromosome 8 (Chr8) gain (77%), consistent with published data. Pre-clinical models that capture this molecular heterogeneity are needed to identify and prioritize effective drug candidates for clinical translation. Here, we describe the successful development of a medium-throughput *ex vivo* 3D microtissue model with several advantages over 2D cell line growth, which can be utilized to predict drug response *in vivo*. Herein, we present proof-of-principle of this PDX-to-microtissue system, using four genomically representative MPNST and three drugs. This work highlights the development of a novel *ex vivo* to *in vivo* preclinical platform in MPNST that successfully captures the genomic diversity observed in patients and represents a resource to identify future therapeutic strategies.

## Introduction

Malignant peripheral nerve sheath tumors (or MPNST) are a heterogeneous group of highly aggressive soft tissue sarcomas with limited treatment strategies. These sarcomas have high recurrence rates and most patients die within five years of diagnosis (Evans, 2002; Hruban *et al*, 1990; Wong *et al*, 1998; Kourea *et al*, 1998; Reilly *et al*, 2017). *NF1* gene inactivation and loss of NF1 protein (neurofibromin) expression characterize the majority of NF1-MPNST (Prudner *et al*, 2020). While *NF1* loss is necessary for MPNST development, it is not sufficient for malignant transformation (Zheng *et al*, 2008; Yang *et al*, 2008; Zhu *et al*, 2002). The cooperating secondary genetic alterations that promote MPNST development include loss-of-function (LOF) alterations in *TP53, CDKN2A*, and polycomb repressor complex 2 (PRC2) complex genes (*EED/SUZ12*) (Baude *et al*, 2014, 2; Lee *et al*, 2014; Cichowski *et al*, 1999; Legius *et al*, 1994; Perry *et al*, 2002; Inoue *et al*, 2021; Kochat *et al*, 2021; Pemov *et al*, 2019). It is increasingly clear that in addition to the common LOF events, a large spectrum of other genetic alterations are present in MPNST including rearrangements in Chr8 and other copy number changes (Dehner *et al*, 2021; Keng *et al*, 2012; Rahrmann *et al*, 2013; Vélez-Reyes *et al*, 2021). One of the major barriers to improving outcomes for MPNST patients is the absence of preclinical models that accurately represent this genetic heterogeneity. To address this issue, we have generated the largest set of genomically characterized NF1-MPNST patient derived xenografts (PDX) (Dehner *et al*, 2021). We have previously shown that these lines harbor the spectrum of genomic alterations that are seen in patients with NF1, including germline and somatic *NF1* mutations, as well as loss of *CDKN2A, TP53* mutations, *EED/SUZ12* mutations, and numerous copy number changes (Dehner *et al*, 2021), and thus may serve as the ideal platform for preclinical discovery. The motivation for generating and utilizing these PDX models instead of traditional cell lines is guided by strong scientific evidence that PDX provide a more genomically-authentic and patient-relevant model for pre-clinical drug evaluation than established cell lines (Koga & Ochiai, 2019).

Human disease biology is profoundly complex and certain biological processes cannot be reproduced on plastic. Additionally, 2-dimensional (2D) cell cultures are prone to genetic drift and monolayer cultures are not representative of human disease (Golan *et al*, 2018; Tignanelli *et al*, 2015; Li *et al*, 2013). In that respect, 3-dimensional (3D) cultures bridge the gap between human cells and animal-based models (Chiovaro *et al*, 2019; Kim *et al*, 2020). These cultures mimic *in vivo* intercellular interactions in 3D in a medium-to high-throughput manner and allow the study of complex tissue structures. Further, it has been well-established that 3D cultures are more predictive of clinical drug response than 2D cultures (Weigelt *et al*, 2010; Jacobi *et al*, 2017). Herein, we have cultured PDX cells *ex vivo* in engineered 3D microenvironments. This provides the opportunity to recapitulate critical cell-cell and cell-matrix interactions that influence drug responses in human tumors that are not taken into account in other models. 3D culture models have been extensively used for cancer drug screening studies, but limited studies exist for NF1-MPNST (Roy *et al*, 2020; Moll *et al*, 2013; Oyama *et al*, 2020).

In this study, we report the successful development of a novel 3D collagen/Matrigel microenvironment (hereafter referred to as microtissue) platform to assess drug response in MPNST. The cells in our 3D microtissue model grow in association with a dense extracellular matrix (ECM), and hence, act more like a developing tumor rather than a monolayer cell culture. Consequently, we expect the drug responses in 3D microtissues to be more representative of the *in vivo* drug response than what could be achieved in 2D culture. We find that PDX which do not form 3D spheres in liquid culture, and PDX from which permanent cell lines could not be derived, were suitable for drug testing in our 3D microtissue platform. Our comparison of *ex vivo* microtissue drug response and *in vivo* PDX tumor growth data revealed significant similarities in drug responses between the two model systems. Taken together, this *ex vivo* PDX-microtissue to *in vivo* PDX platform offers an ideal model system for drug discovery, drug response validation, and exploration of MPNST biology in a genomically heterogeneous system that is representative of the human condition.

## Results

### PDX established from human MPNST represent full tumoral heterogeneity

PDX serve as a useful preclinical model for cancer drug screening studies. We previously reported the establishment of eight MPNST PDX lines from patient tumors (Dehner *et al*, 2021). These were established by engrafting minced or single cell suspended tumor into immunodeficient NOD.Cg-*Prkdc*^*scid*^ *Il2rg*^*tm1Wjl*^/SzJ (NSG) mice and serially passaging them through additional mice (**Fig 1A**). Here, we broaden our sample size by including five additional PDX-tumor pairs. Five MPNST PDX lines were established from biopsy-proven NF1-MPNST between 2014 and 2021 at three institutions: Washington University, Johns Hopkins University, and University of Minnesota. Clinical parameters from the thirteen patients are summarized in **Table 1**. Eight of the patients were male, with a median age of onset of 30.5 years (range, 9–67 years).

**Table 1.** MPNST PDX lines and data collected to date.

| Sample | Age group (years) | Sex | Microtissue Quality | MPNST | Location | Size (cm) | Clinical Status |
| --- | --- | --- | --- | --- | --- | --- | --- |
| WU-225 | 18-40 | Male | Good | Primary | Thigh | >10 | Deceased |
| WU-356 | 18-40 | Male | Unusable | Primary | Mediastinum | >10 | Deceased |
| WU-368 | >40 | Female | Unusable | Primary | Calf | >10 | NED |
| WU-386 | 18-40 | Male | Unusable | Primary | Neck | <10 | NED |
| WU-436 | 18-40 | Male | Unusable | Primary | Thigh | >10 | NED |
| WU-487 | >40 | Male | ND | Recurrent | Left Neck | <10 | Deceased |
| WU-561 | 18-40 | Female | ND | Recurrent | Left Pelvis | >10 | Alive with metastatic disease |
| JH-2-002 | <10 | Male | Robust | Primary | Pelvis | <10 | NED |
| JH-2-023 | 18-40 | Male | ND | Primary | Paraspinal | <10 | NED |
| JH-2-031 | 10-17 | Male | Unusable | Primary | Retroperitoneal | >10 | Deceased |
| JH-2-055-b | 10-17 | Female | ND | Primary | Scalp/Neck | <10 | Alive with metastatic disease |
| JH-2-079-c | 10-17 | Female | Good | Recurrent | Thigh | <10 | Deceased |
| MN-2 | >40 | Female | Robust | Primary | Maxillary Sinus | <10 | Unknown |

*ND = not determined. NED = no evidence of disease.*

**Figure 1.**
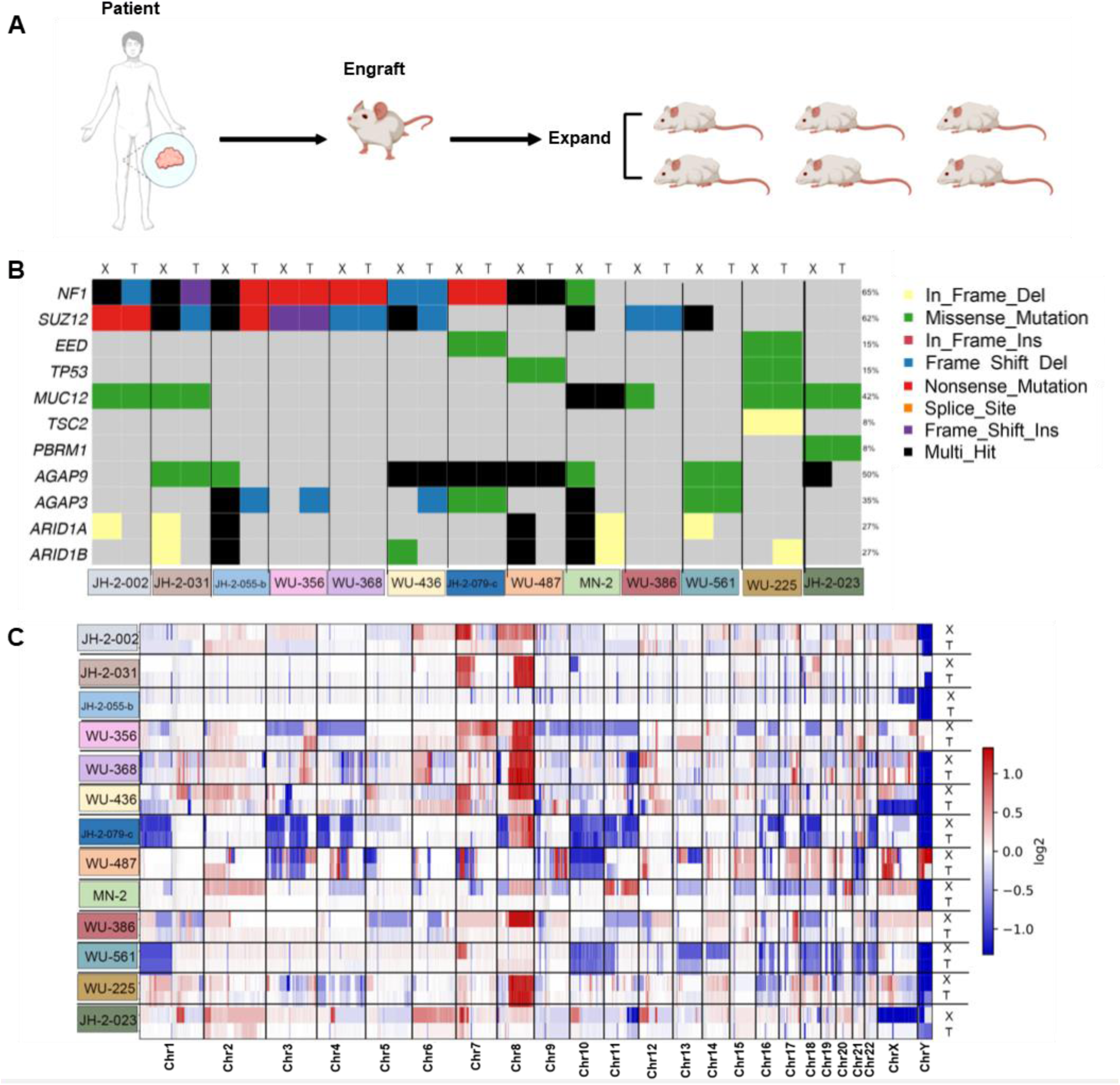
Diverse MPNST patient tumors used in separate models. A. Schematic description of patient derived xenografts engrafted into NRG mice for drug studies. B. Heatmap of single nucleotide variants across all 13 PDX-tumor pairs (X denotes PDX; T denotes parental tumor). Each somatic variant is present in only a fraction of the samples. Percent-wise distribution is shown on the right. C. Copy number variations in all 13 PDX-tumor pairs.

To determine the inter-tumoral heterogeneity across the 13 PDX-MPNST pairs, we performed deep whole exome sequencing to assess single nucleotide variants (**Fig 1C**) and whole genome sequencing for copy number alteration calling (**Fig 1D**). Consistent with previous studies, we observed some of the common mutations associated with MPNST, including *SUZ12* (7 of 13 pairs), *TP53* (2 of 13 pairs), and *EED* (2 of 13 pairs) (**Fig 1C**). Some unique mutations were also observed in our analysis, including *MUC12* (3 of 13 pairs), *AGAP3* (3 of 13 pairs), and *AGAP9* (5 of 13 pairs). MUC12 is a membrane glycoprotein that is reported to be a biomarker and metastasis promoter for stage II and stage III colorectal cancer (Matsuyama *et al*, 2010, 12); the AGAP family of proteins have also been reported to promote cancer cell invasiveness (Yoo *et al*, 2020). Mutations in the DNA repair gene, *PBRM1* were observed in 1 of 13 pairs. *PBRM1* (polybromo1) is a DNA binding protein that is required for cell cycle progression through mitosis. *PBRM1* mutant patients have been reported to respond poorly to immunotherapy (Yang *et al*, 2021). Copy number analysis also revealed significant genomic heterogeneity, with Chr8 aneuploidy being the most common event (10 of 13 pairs; **Fig 1D**). Notably, some genes were found to have somatic variants only in the PDX but not in the parental tumors. This finding can be attributed to expansion of select cell populations in mice during PDX establishment, or sampling bias due to intra-tumoral heterogeneity. As a consequence, certain variants are observed at a higher frequency in PDX than the parental tumors, which is suggestive of engraftment-related clonal selection within PDX, as observed previously (Peille *et al*, 2020). For example, *NF1 and SUZ12* were mutated only in the PDX line of MN-2, but not in the parental tumor. We believe that an excess of non/pre-tumorigenic cells in the MN-2 tumor and clonal outgrowth in the corresponding PDX contributed to this genetic variability.

### Dissociated PDX exhibit limited growth in two-dimensional culture and as spheroids

Since PDX are a costly and time-consuming model for *in vivo* drug studies, we were interested in developing an alternative platform that maintains MPNST heterogeneity and is suitable for drug response studies. For the studies described here, PDXs were passaged in NOD-*Rag1*^*null*^ *IL2rg*^*null*^ (NRG) mice. We dissociated PDX samples from mice and attempted two-dimensional (2D) tissue culture on plastic. Only six out of the thirteen PDX tested could be established as cell lines, while others did not grow more than a few days and were not an option for drug studies. Our observations suggest that MPNST cell lines develop at a low frequency, thereby highlighting the need for an alternative, genomically heterogenous model system. Further, the majority of the PDX cells could not form spheroids in a collagen-free environment. After two days of growth in the collagen-free format, MN-2 and JH-2-031 began to aggregate while cells from many other samples remained dispersed (**Fig EV1A**). After seven days in culture, MN-2 exhibited a tight spheroid formation while JH-2-031 cells remained a loose aggregate (**Fig EV1B**). The limited growth in 2D culture and as spheroids suggest that these applications of PDX are not viable options for drug response studies. Additionally, neither of these two culture conditions reflect a developing tumor as they lack sufficient stroma and components of the ECM.

### Three-dimensional engineered microenvironments, or 3D microtissues, are successfully created from dissociated PDX with notable differences

After failing to create reliable 2D or spheroid cultures from PDX, we attempted to create three-dimensional (3D) microtissues, based on prior experiences with other cell types (Crampton *et al*, 2018). Dissociated PDX were depleted of mouse and dead cells and assembled into 3D microtissues with the addition of collagen and Matrigel for tissue support (full details in Methods) (**Fig 2A**). We were able to capture a high percentage of viable human tumor cells from PDX samples and use them efficiently in the microtissue platform. Unsurprisingly, variation was observed for each dissociated PDX that formed a 3D microtissue. In the 3D microtissue format, some PDX failed to proliferate or grew only for a few days, and thus, were removed from drug studies (**Fig EV2**). We also observed a few PDX remodeled their collagen matrix. As quickly as two days in 3D microtissue culture, MN-2 showed signs of compacting its collagen fibers (**Fig EV3**). After seven days, MN-2 completely remodeled its microenvironment forming a small round mass of cells and collagen. Other PDX took longer to remodel their microenvironment (11 days for JH-2-079-c) or not at all (as with WU-225; **Fig EV3**). Differences in collagen compaction rates have been observed previously (Crampton *et al*, 2019).

**Figure 2.**
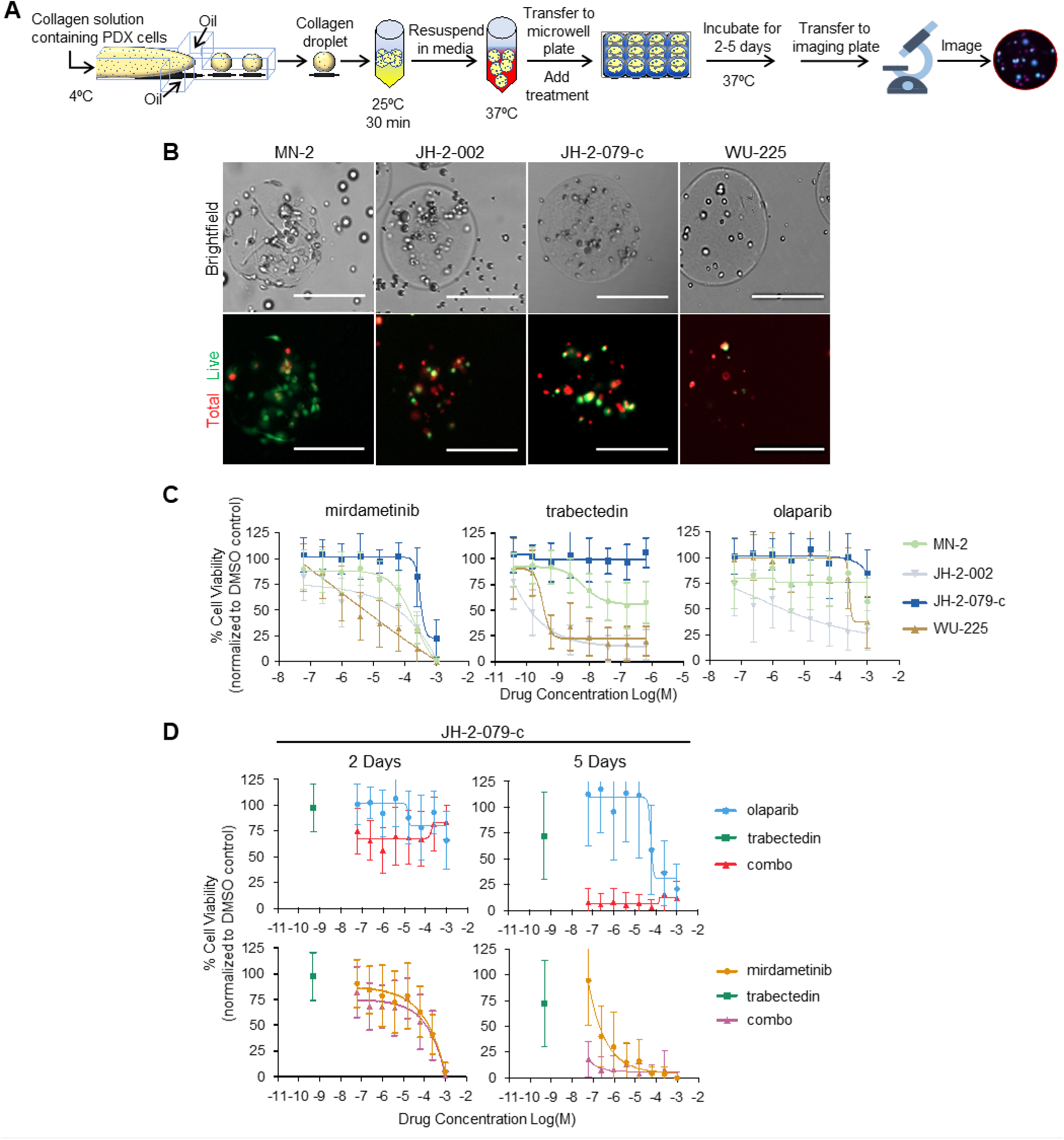
PDX 3D microtissues are inhibited with trabectedin drug combinations. A. Schematic description of dissociated patient derived xenograft tumors processed into 3D microtissues. B. Representative brightfield and fluorescent images of PDX 3D microtissues made from MN-2, JH-2-002, JH-2-079-c and WU-225 stained with calcein AM (live) and DRAQ5 (total) after two days in culture. Scale bars: 200μm. C. Dose response curves of four PDX 3D microtissues exposed to single agents mirdametinib, trabectedin or olaparib for two days in culture. Data points and error bars represent mean±SD, n≥10. D. Dose response curves of JH-2-079-c exposed to trabectedin combinations with either mirdametinib or olaparib for two or five days in 3D microtissues. Trabectedin concentration was kept constant (0.5 μM) with varied concentrations of mirdmetinib or olaparib. Data points and error bars represent mean±SD, n≥10.

Given the variability, 3D microtissues were ranked for quality based on the initial percent cell viability and the difference between percent cell viability at 48 hours after assembly and just prior to assembly. We qualified them as robust (>90% viability), good (>50%), or unusable (<50%) and only used those categorized as robust or good for drug studies. Four out of nine PDX-microtissues tested met these criteria - MN-2, JH-2-002, JH-2-079-c and WU-225 (**Table 1**). Here we show the successful development of a 3D microtissue platform generated from PDX (**Fig 2B**), offering a more biologically relevant model than 2D or spheroid culture.

### Initial microtissue drug studies demonstrate selective response to single agents and combination therapies

In order to test the feasibility of this novel 3D platform as a drug screening model, we chose three drugs as proof of principle. We chose drugs predicted to have a response based on mutations present in our PDX-tumor pairs. Prior studies showed mutations in DNA repair genes in up to 25% of MPNST ((Dehner *et al*, 2021; Kaplan *et al*, 2018) (**Fig 1C**), leading to selection of olaparib, a PARP inhibitor, for our initial studies. Additionally, we selected trabectedin as a chemotherapeutic that could be combined with a PARP inhibitor or other targeted therapies. Trabectedin works by distorting the structure of DNA and interfering with the nucleotide excision repair (NER) system (Dubois & Cohen, 2009), thus potentially leading to synthetic lethality in combination with a PARP inhibitor. In fact, olaparib and trabectedin have already been used in combination in a Phase 2b trial to treat advanced soft-tissue sarcomas (Grignani *et al*, 2018). Finally, we chose mirdametinib (previously known as PD0325901) (Barrett *et al*, 2008) as a representative MEK inhibitor, as *NF1* loss results in hyperactive RAS-RAF-MEK-ERK signaling and MEK inhibition has previously been shown to be partially active in NF1-associated tumor cells (Jousma *et al*, 2015; Weiss *et al*, 2021; Dodd *et al*, 2013; LoRusso *et al*, 2010).

Since 3D microtissues generated from PDX are more reflective of the human disease than 2D or spheroid culture, we hypothesized that drug response studies in our 3D microtissues would inform *in vivo* activity. We initially tested trabectedin plus mirdametinib and trabectedin plus olaparib with both combinations in a fixed ratio of 1:2000 at each dose. We tested each drug with an 8-point dose range with 4-fold dilutions between each dose to achieve a wide range of concentrations. Each PDX 3D microtissue was treated with drug or vehicle control for different experimental time points. Mirdametinib had the strongest effect on cell viability for each of the four microtissues tested, while olaparib or trabectedin had differential effects depending on the PDX (**Fig 2C**). Notably, JH-2-079-c responded the least to all three drugs. This PDX is also the only PDX derived from recurrent tumor that was tested in our 3D microtissue drug studies and thus, may be a more drug-resistant tumor. WU-225 was noted to be the most responsive to trabectedin and mirdametinib (**Fig 2C, Fig EV4A**). While both WU-225 and JH-2-079-c have mutations in components of the PRC2 complex, WU-225 has additional missense mutations in *TP53* (**Fig 1B**).

### Low concentration of trabectedin and longer exposure time enhances cytotoxicity in 3D microtissue combination studies

With our initial fixed ratio, drug combinations did not show an enhanced effect over single agents in any of the four PDX 3D microtissues (**Fig EV4A & B**). We hypothesized we were likely missing any synergistic or additive effect in the fixed ratio regimen due to both the short time course and the robust response to high doses of trabectedin alone. Therefore, we chose a low dose of trabectedin (0.5 nM) and maintained this concentration for each of the eight doses of either olaparib or mirdametinib. An enhanced effect emerged with this new regimen in both combinations, but was more pronounced with olaparib plus trabectedin (**Fig 2D**). Additionally, JH-2-079-c exhibited a dramatic reduction in cell viability after five days in culture, even at the lowest concentrations of olaparib or mirdametinib (**Fig 2D**). These data show that drug synergy can be observed in NF1-MPNST PDX cells cultured in 3D microtissues cultured for longer than 2 days.

### Trabectedin treatment inhibits *in vivo* tumor growth in PDX models of MPNST, which corroborates the findings from our 3D microtissue model

Next, we sought to validate our findings from microtissues in our *in vivo* PDX model. Tumor bearing NOD-*Rag1*^*null*^ *IL2rg*^*null*^ (NRG) mice were administered drugs either as single agents or in combination as shown in **Fig 3A**. MN-2 showed modest sensitivity to trabectedin and a more potent anti-tumor effect from the combination of trabectedin and mirdametinib (**Fig 3B**). These drugs had similar single agent responses in the microtissue model. In contrast, WU-225 demonstrated enhanced tumor growth inhibition to single agent trabectedin (**Fig 3B**). Dramatic inhibition in tumor growth was observed for WU-225 when treated with a combination of trabectedin and mirdametinib or trabectedin and olaparib (**Fig 3B, C**). Again, these drugs had single agent activity in the *ex vivo* microtissue model. Interestingly, JH-2-079-c had limited response to single agents *in vivo*, similar to what was seen *ex vivo*. However, combination therapy with trabectedin and olaparib (**Fig 3C**, p=0.0164) and trabectedin and mirdametinib (**Fig 3B**, P < 0.05) did significantly decrease tumor volume *in vivo*, similar to what was seen with the five-day *ex vivo* studies. In contrast, MN-2 demonstrated enhanced sensitivity to trabectedin alone (P < 0.0001) as well as mirdametinib single agent (P < 0.05). However, no increase in effectiveness was observed with trabectedin and olaparib combination therapy, while a significant benefit was observed with the combination of trabectedin and mirdametinib versus trabectedin treatment alone (**Fig 3B, C**). Taken together, these data confirm that the findings from the 3D microtissue model are predictive of *in vivo* response.

**Figure 3.**
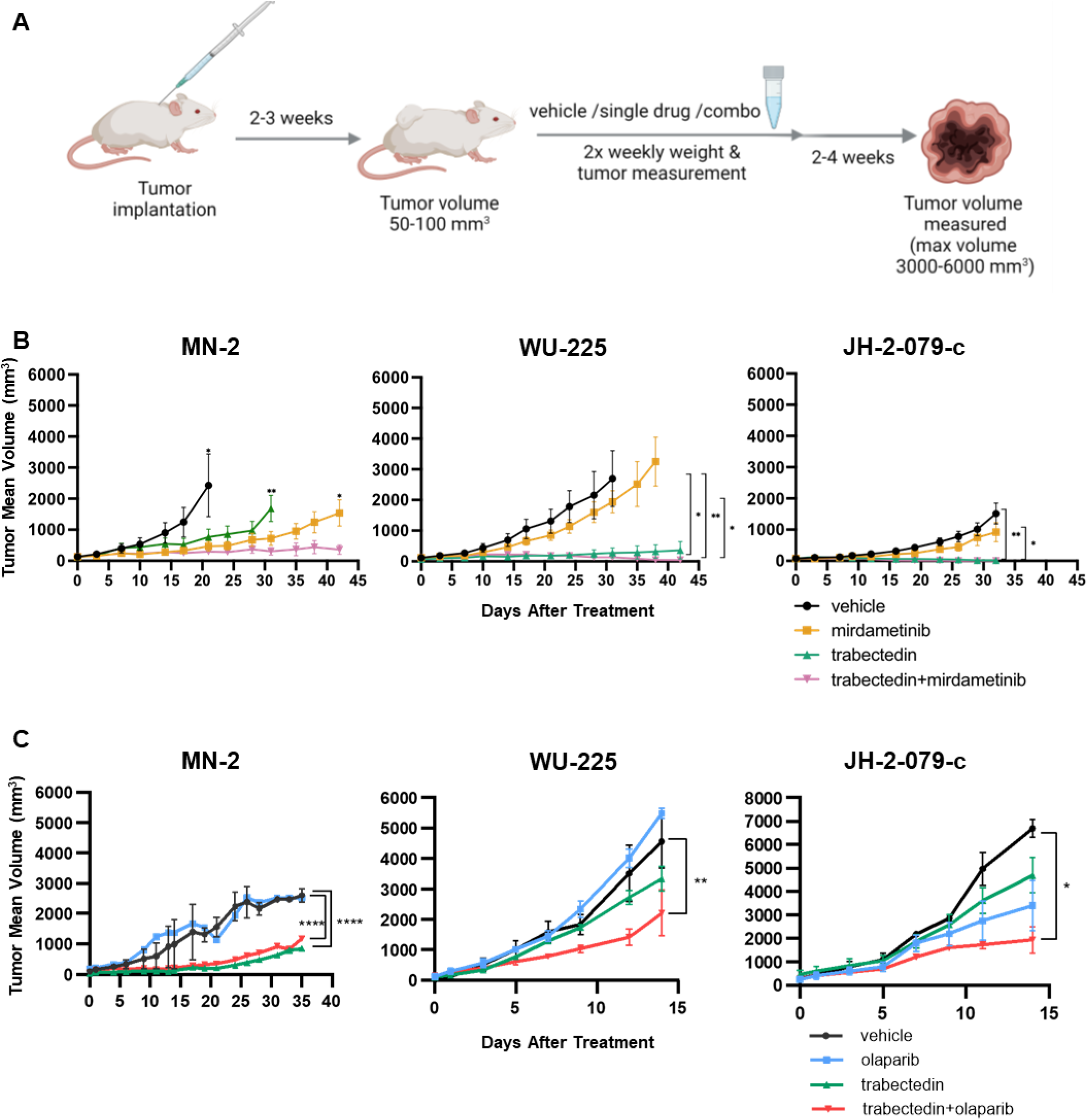
*In vivo* tumor growth is inhibited with trabectedin treatments. A. Schema describing timeline of drug administration following PDX engraftment. B. Tumor volume response to mirdametinib, trabectedin or the combination of the two in MN-2, WU-225, and JH-2-079-c PDX grown in mice. C. Tumor volume response to olaparib, trabectedin or the combination of the two in MN-2, WU-225, and JH-2-079-c PDXs grown in mice. Data points and error bars represent mean±SEM n=3-5, ANOVA was used to assess statistical significance (****P < 0.0001; ***P < 0.001; **P < 0.01; *P < 0.05).

### Gene expression analysis of PDX samples identifies key pathways that may predict the quality of an assembled 3D microtissue

Given the genetic diversity of the PDX samples as well as the phenotypic differences in their ability to grow within the microtissue framework, we evaluated gene expression signatures of PDX cells by RNA-sequencing, as depicted in **Fig 4A**. Mapping the gene expression of each sample using two principal components, we observed that the first principal component, depicted on the x-axis, segregates most of the samples that grow well in the microtissues (left) from those that do not grow well (right). Such clustering pattern suggests that we can predict the growth characteristics of 3D microtissues within our platform on the basis of their gene expression data. Our observations have significant implications for future studies as we continue the development of additional 3D microtissues. WU-225, however, appears to be an outlier in **Fig 4A**.

**Figure 4.**
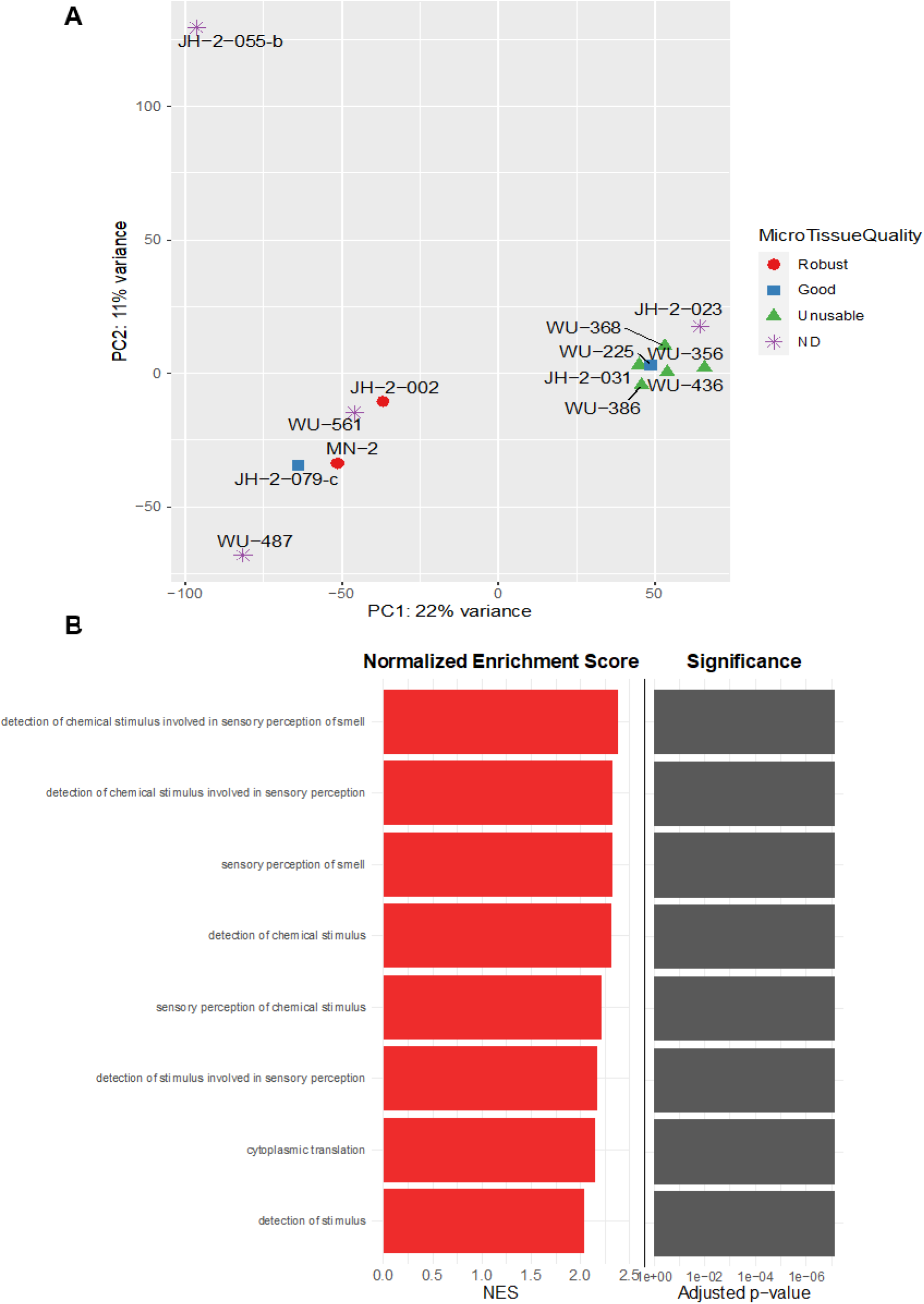
Gene expression analysis of microtissues by their quality. A. Two-dimensional embedding of gene expression of each patient sample. Shape indicates microtissue quality from Table 1. B. Biological processes enriched in microtissues that exhibit robust or good growth compared to those that do not grow in culture. Adjusted p-value < 0.01.

We attempted to identify whether there were specific biological pathways that were differentially represented between the two sets of samples. We found that 525 genes were differentially expressed (adjusted p < 0.01) (**Table EV1**). Gene enrichment analysis identified pathways related to perception of external (chemical) stimuli to be enriched in “robust” and “good” microtissues (**Fig 4B, Fig EV5**), which is correlated with their enhanced cellular ability to respond to signals in their extracellular environment (Rodrigues *et al*, 2021; Wan *et al*, 2017).

## Discussion

Mutational heterogeneity and DNA aneuploidy commonly seen in MPNST present challenges to the successful development of effective therapies for patients with MPNST. Many pre-clinical platforms rely on genetically engineered mouse models which are not capable of representing the full heterogeneity of genomic alterations that exist in patient tumors, limiting the translatability of pre-clinical studies to the clinic. To address this problem, we have developed a PDX-based *ex vivo* 3D microtissue platform that we have described herein. The entire system offers several advantages, such as the ability to perform rapid and cost-effective drug response studies, the capacity to validate drug effectiveness, and the potential to alter the tumor microenvironment by addition or removal of biologically relevant ECM components, or inclusion of additional tumor stromal cell types.

Our PDX tumor model successfully reflects the genomic diversity of MPNST and most PDX accurately resemble the genetic signature of their parental tumor. We identified the variety of genetic alterations that are typical of NF1-MPNST in the 13 PDX-MPNST pairs. We identified somatic *NF1* mutations in 8 of 13 pairs. Five cases, WU-386, WU-561, WU-225, MN-2, and JH-2-023, were presumed to have a second hit in *NF1* through another mechanism. Prior studies also report the occurrence of NF1 cases without an identifiable second hit in *NF1* (Gosline *et al*, 2017); these cases presumably have mutation(s) in non-coding region(s) or copy number loss of *NF1. SUZ12* and *EED* together form a part of the PRC2 complex that is commonly mutated in MPNST, particularly in combination with *NF1* mutations (Lee *et al*, 2014, 2; Zhang *et al*, 2014). We also observed mutations in both *NF1* and *SUZ12*/*EED* in 7 of 13 PDX-MPNST pairs. *TP53*, another common genetic variant seen in MPNST, was mutated in 2 of 13 pairs. We additionally show the presence of somatic variants not yet reported in MPNST – *AGAP3/9*, and *MUC12*. While mutations in *AGAP3/9*, and *MUC12* have been associated with inferior disease prognosis in other cancers (Matsuyama *et al*, 2010, 12; Yoo *et al*, 2020), none have been reported to be altered in soft tissue sarcomas to date. While our PDX are overall representative of the parental tumors, some degree of clonal evolution was observed in PDX lines for WU-386, JH-2-055-b, and JH-2-002, where mutations not seen in the parental tumor were observed in the PDX. This finding is in line with other studies (Richter-Pechańska *et al*, 2018) in which authors observed that clonal complexity of parental leukemia may be altered in the corresponding PDX. Variations between primary tumor and PDX samples are likely indicative of clonal selection, where the clone responsible for the divergence may already be present at a subclonal level in the parental tumor sample.

Our PDX model can be utilized in a medium-to high-throughput manner for preclinical studies. While the *in vivo* PDX model has strong advantages for drug efficacy and sensitivity studies, experiments using them can be both time-intensive and costly. In the last decade, engineered 3D microenvironments have garnered considerable attention as an alternative preclinical model to study cancer cells. These cellular models are small formations that closely mimic the natural tumor microenvironment without the complexity and cost of maintaining animal colonies needed for *in vivo* studies (Brancato *et al*, 2017, 2018). As sensitivity of cancer cells to drug treatment is strongly influenced by the tumor stroma (Brancato *et al*, 2020; Landry *et al*, 2018), the 3D microtissues developed from our PDX lines are an ideal platform to study drug response of MPNST. The collagen matrix used in our 3D microtissues, which is absent in traditional 2D culture, favors cell-cell interactions by mimicking the ECM (Brett *et al*, 2016; Crampton *et al*, 2018). Additionally, we found that dissociated PDX tumor cells could rarely grow as tight spheroids, rendering this 3D model as less than ideal.

Utilization of our system has already yielded important information regarding the potential activity of some therapeutic agents. Assessment of cell viability following treatment with a select set of drugs identified mirdametinib as a potent small molecule therapeutic agent against all microtissues tested. Mixed response was observed for trabectedin or olaparib depending on the PDX. Both WU-225 and JH-2-002 have mutated PRC2 components (*EED* and *SUZ12*, respectively) which could explain their uniquely similar response curves to trabectedin and mirdametinib. JH-2-079-c, however, did not respond to either single agent trabectedin or olaparib after two days in culture. We believe this differential response is due to the inherent genetic variability of these MPNST. JH-2-079-c is the only MPNST in our sample set with mutations in all three genes - *EED, AGAP3*, and *AGAP9*. Additionally, this line was generated from a recurrent tumor in a patient who had previously received chemotherapy for the initial MPNST, and thus, may be inherently more aggressive and drug resistant. Our data indicate a possible role of the somatically acquired genetic variability of MPNST in determining drug response.

We observed similar response to single agents in both the 3D microtissue and *in vivo* PDX models. While synergy was not observed in our initial 1:2000 combination 3D microtissue studies, an enhanced drug effect was observed when we maintained a low trabectedin dose in combination with olaparib or mirdametinib for two days. Additionally, this effect was enhanced when JH-2-079-c was grown longer for five days as a 3D microtissue. This extended drug exposure of 3D microtissues better reflects an *in vivo* model where animals are treated for weeks. Indeed, when grown for more than two days, the drug response of three therapeutic drugs and their combinations in 3D microtissues was validated in our *in vivo* PDX model.

Finally, our 3D microtissue system offers a chance to identify different factors that may be important for drug response. This system will allow other factors such as ECM components and other cell types to be evaluated in isolation as has been done in organoid systems for other cancer types. For example, Raghavan et al (Raghavan *et al*, 2021) report differential response of patient-matched organoids to chemotherapeutic agents against PDAC (pancreatic ductal adenocarcinoma). Addition/removal of paracrine factors from organoid growth media was able to significantly modulate the drug response pattern of these *ex vivo* models. Such paracrine factors, along-with the non-malignant cells present within metastatic sites, make up the tumor microenvironment (TME) that holds the potential to shape cancer cell state. Hence, it is possible that the microtissue model may also be enhanced by the addition of specific cytokines and/or conditioned growth media derived from other MPNST cell lines.

In accordance with Raghavan et al (Raghavan *et al*, 2021), the observed drug response pattern of our 3D microtissue model may be linked to differences in gene expression. We observed homologous transcriptional clustering across microtissues that were categorized as either “robust” or “good” versus “unusable”. Pathway enrichment analysis identified genes related to perception of external stimuli in these microtissues, which may be linked to the ability of these lines to grow well. Rodrigues et al (Rodrigues *et al*, 2021) also report that cancer cells grown in 3D cultures are more receptive to environmental stimuli from all directions that properly represents the *in vivo* stimuli pattern. Further, Wan et al (Wan *et al*, 2017) report the development of 3D system for cancer cells that promotes “natural” tumor growth in response to chemical stimuli in a controlled 3D environment. Hence, enhanced receptiveness to environmental stimuli might be crucial for the development of viable 3D microtissues.

The notable limitations of our PDX-to-microtissue model are the low success rate of establishing “robust” or “good” microtissues, and limited growth duration of established 3D microtissues. We expect that modification of growth media components will enhance the durability of our novel platform. Efforts are underway to improve growth conditions that permit the establishment of additional 3D microtissues that can grow for longer periods of time. Further, improved methodology that allows us to evaluate viable cell numbers is under optimization.

In summary, we report the development of a novel model system for testing MPNST response to candidate drugs. The combination of an *ex vivo* microtissue platform followed by *in vivo* PDX testing allows for medium-to high-throughput drug screening and *in vivo* validation. Future studies will be aimed at testing new combinations of drugs as well as dissecting out components of the TME which may be important for drug response through modifications to the microtissue culture conditions in terms of cell types and ECM components.

## Materials and Methods

### Study approval

Study material was collected under IRB-approved protocols at Johns Hopkins University (protocol no. J1649) (Pollard *et al*, 2020), Washington University at St. Louis (protocol no. 201203042), and University of Minnesota (protocol no. STUDY00004719). All subjects provided written informed consent prior to participation in the study.

### Human subjects and PDX tumor model establishment

Two to four tumor pieces were collected following surgical removal of patient tumors. The specimens were placed in DMEM containing 10% FBS during transportation to the laboratory and were later used for implantation into mice. Tumor tissue was implanted dorsally into 5-to 6-week-old NOD-*Rag1*^*null*^ *IL2rg*^*null*^ (NRG) mice and allowed to grow. Once the tumors were ∼2cm x 2cm in size (or the mouse met other parameters that required its sacrifice), tumors were removed, minced, and engrafted into additional mice. This process was repeated for six passages. Engraftment success was defined as the ability of the PDX line to be serially transplanted for six passages. JH-2-055-b represents the same line previously published as JH-2-055, but multiple samples have been processed from a single patient, and therefore JH-2-055-b has been renamed accordingly (Wang *et al*, 2020).

### WES, WGS, and RNA-seq library preparation and sequencing

Normal germline samples, tumor samples, and xenograft samples from six patients (total 18 samples) were used for DNA sequencing. DNA library was constructed using KAPA HyperPrep Kits for NGS DNA Library Prep. For whole exome sequencing, exomes were captured by IDT exome reagentxGen Exome Research Panel V1.0. Total of 18 exome libraries were sequenced by NovaSeq6000 S4 300XP with ∼200× coverage for normal samples and 800-1000x coverage for tumor/xenograft samples. For WGS, a total of 16 libraries (two normal samples were not sequenced) were sequenced by NovaSeq6000 S4 300XP with 15-20x coverage. For RNA-seq, tumor and xenograft samples were used from six patients. Samples were prepared using TrueSeq stranded total RNA library kit with Ribo-Zero for rRNA depletion. Libraries were sequenced by NovaSeq6000 S4 300XP with targeted coverage of 30M reads per sample.

### WES data analysis

WES data from eight patients reported in our previous paper (Dehner *et al*, 2021) were downloaded from the NF Data Portal. WES sequencing FASTQ files were trimmed using Trimmomatic v 0.39 (Bolger *et al*, 2014) and aligned against reference sequence hg38 via BWA-MEM (Li, 2013). Duplicated reads were marked using picard “Markduplicates” function. GATK V4.2 base quality score recalibration (BQSR) was also used to process BAM files. For PDX sequence data, Xenosplit was used to filter out mouse-derived reads using mouse (GRCm38) and human (hg38) reference genomes. Somatic single nucleotide variants (SNVs) and small insertions or deletions (indels) were detected using VarScan2 (Koboldt *et al*, 2012), Strelka2 (Kim *et al*, 2018), MuTect2 (Benjamin *et al*, 2019), and Pindel (Ye *et al*, 2009). Variant filtering and annotation were done using Variant Effect Predictor (VEP) (McLaren *et al*, 2016). Common variants found in the 1000 Genomes MAF and GnomAD MAF > 0.05 were filtered out. One more step was added to the tumor sample variant calling by checking SNV/indels from paired xenograft in the sample because some variants were missing in the tumor samples. This was because the tumor purity was lower than xenograft sample purity (Xenograft sample has >90% purity). If new SNV/indels from paired xenograft were found in the tumor sample, these variants would be added to the tumor variant calling result. Waterfall somatic variant plots were created with maftools v2.10.0 (Mayakonda *et al*, 2018).

### Copy number variant (CNV) analysis using WGS data

WGS data from eight patients reported in our previous paper (Dehner *et al*, 2021) were downloaded from Synapse. Alignment of sequence reads, removal of duplicate reads, and BQSR steps are the same as in WES. CNVkit V2.0 was used to infer and visualize copy number. Normal pooled reference was first built from all normal samples. Then, the reference was used to extract copy number information from tumor/xenograft sample BAM files. Heatmap was drawn using CNVkit’s heatmap function.

### Bulk RNA-seq analysis

Tumor and xenograft samples from six patients were used for RNAseq. RNA was prepared with a TrueSeq stranded total RNA library kit with the removal of rRNA using Ribo-Zero. The libraries were sequenced by NovaSeq6000 S4 300XP with 30M reads. Aligned RNA reads were aligned to their respective genome assembly. Initial primary tumors and PDX samples were aligned to GRCh37. Mouse-derived reads were filtered using Xenosplit. RNA reads were quantified using the Salmon algorithm (Patro *et al*, 2017). Gene counts and transcript counts were normalized by using the DESeq2 package (Love *et al*, 2014, 2).

### PDX Cell Dissociation for 3D microtissue assembly

All mouse experiments were approved by the Institutional Animal Care and Use Committee at University of Minnesota under protocol #2101-38758A. For each PDX tumor, a tissue sample was minced in a basement membrane matrix and passed through a 1mL syringe with an 18-gauge needle to achieve a homogenous mixture, which was injected into the flank of NRG mice (Jackson Labs). Each PDX tissue sample was sufficient to implant two mice. Enough human PDX was used to implant 5 animals. Once the xenograft tumors reached the maximum size allowed (2000mm^3^), mice were euthanized, and the tumors extracted in a laminar flow hood under sterile conditions. The xenograft tumors were immediately digested using the human Tumor Dissociation Kit (130-095-929, Miltenyi Biotec) in combination with the GentleMACS Octo Dissociator (Miltenyi Biotec). The digested PDX was filtered through a 70μm cell strainer and then depleted of residual red blood cells using 1X RBC Lysis Buffer (eBioscience). Dead cells were removed from the digested PDX using a Dead Cell Removal Kit (130-090-101, Miltenyi Biotec). Murine cells were removed from the dead cell depleted fraction using a Mouse Cell Depletion Kit (130-104-694, Miltenyi Biotec). Cell viability was determined using flow cytometry and 7-AAD Viability Staining Solution (Biolegend). Cell counts for all fractions were ascertained using the Countess cell counter (Invitrogen).

### 2D Culture Details

PDX cells were maintained in Dulbecco’s Modified Eagle Medium High Glucose (ThermoFisher), 10% Fetal Bovine Serum (Gibco), and 1% Penicillin-Streptomycin (Gibco). All cultures were maintained at 37°C, 5% CO_2_, atmospheric O_2_ and 95% humidity in tissue culture-treated 24-well plates (Corning).

### Microwell Fabrication

Microwell plates were fabricated following previously established methods (Crampton *et al*, 2019). Briefly, tissue culture-treated multi-well plates were coated with a thin layer of 2% agarose and dehydrated. Polydimethylsiloxane (PDMS) stamps featuring a radial pattern of 300μm posts were then placed immediately upon fresh 2% agarose solution that was pipetted into each well. After cooling, stamps were removed gently and well-plates were washed with appropriate culture medium before adding microtissue solution.

### Microtissue Fabrication

For the assembly of collagen microtissues, previously established protocols were followed (Crampton *et al*, 2018; Brett *et al*, 2016; Crampton *et al*, 2019). Briefly, high-concentration rat tail collagen I (Corning) was buffered with 10 × Dulbecco’s phosphate-buffered saline (DPBS), neutralized to pH 7.4, supplemented with 10% Matrigel (Corning), diluted to 6mg/mL concentration, and mixed with cells (6×10^6^ cells/mL). At 4°C, the collagen solution was partitioned into droplets using a flow-focusing microfluidic device. Tissues were collected in a low-retention Eppendorf tube and polymerized for 30 min at 25°C. The oil phase was then removed and the collagen microtissues were resuspended in culture medium.

### Drug Treatment (3D microtissues)

Inhibitors used in this study were: 1 mM mirdametinib (S1036, Selleckchem) and 1 mM olaparib (S1060, Selleckchem) dissolved in DMSO, and 0.5 μM trabectedin (clinical excess from Washington University pharmacy, manufacturer Johnson & Johnson) dissolved in water. Drug stocks were serially diluted for 8-point dose curves with 4-fold dilutions between each dose. Combinations of trabectedin with mirdametinib or olaparib were kept in a fixed 1:2000 ratio, respectively, for each dose, or with trabectedin at a constant concentration of 0.5 nM. PDX cell-laden microtissue cultures were treated with single and dual agents by addition of the diluted agent(s), in DMSO, to their relevant concentrations in the culture medium. The cell encapsulated microtissue solution was then added to the agent-media solution to obtain desired concentration. Control samples were treated with DMSO (Sigma-Aldrich) diluted in culture medium.

### Cell Viability Assay

Microtissues containing encapsulated PDX cells were fabricated and cultured. At the end of the treatment period, tissues were retrieved from the microwells through gentle pipetting and collected in a 1.5mL tube. To assess cell viability, constructs were washed thoroughly with DPBS (Corning) and then incubated in the dark at 21°C for 30 minutes with a staining solution of 5 μM DRAQ5 (Invitrogen) and 5 μM Calcein AM (ThermoFisher). A Zeiss Axio Observer was used to image z-positions at 50 μm intervals over a distance of 250 μm and images were further analyzed using Cellpose (Stringer et al, 2021) and on ImageJ (NIH). For experimental wells, live cell over total cell count ratio was normalized to vehicle-only treated cells to produce percent cell viability. Data were analyzed in Prism software (GraphPad Prism) and dose-response curves generated using nonlinear regression Log(inhibitor) vs. response-variable slope model.

### Drug Treatment (PDX)

All animal experiments were approved by the Institutional Animal Care and Use Committee at Johns Hopkins under protocol #MO19M115 and at Washington University under protocol #20190118. Female NRG mice were purchased from Jackson Laboratory. For trabectedin and olaparib treatments, single cell suspension of tumor cells (3×10^6^ cells per mouse) was implanted subcutaneously in the flank of 6-to 8-week-old NRG mice. For trabectedin and mirdametinib experiments, minced tumor fragments were implanted subcutaneously in the flank of 6-to 8-week-old NRG mice. Drug treatment started when tumor-bearing mice reached approximately 50-100 mm^3^ in volume and commenced for three weeks (trabectedin and olaparib) and 5-6 weeks (trabectedin and mirdametinib). Mice were randomized at 3-5 animals per treatment group to achieve similar mean tumor volume between groups. Drug doses and modes of delivery used in this study were: trabectedin (clinical access from Washington University pharmacy, manufacturer Johnson & Johnson) at 0.15 mg/kg weekly via tail vein injection; mirdametinib (SpringWorks Therapeutics, formulated in 0.5% HPMC and 0.2% Tween 80) at 1.5 mg/kg via oral gavage daily; and olaparib (Selleckchem) at 100 mg/kg via oral gavage daily. Tumors were monitored twice (Johns Hopkins) and thrice (Washington University) weekly with caliper measurements and tumor volume was calculated using the formula L x W^2^(π/6), where L is the longest diameter and W is the width. Animals were euthanized when they reached ∼2 cm in any dimension or reached other ethical endpoints as defined by the institution’s animal protocol.

### Statistics

Two-way ANOVA was used to calculate statistical significance. Analyses were considered statistically significant if P < 0.05.

## Data availability

Data (*in vitro* and *in vivo* drug studies, RNA sequencing of PDX cells and genomics) generated in this study has been deposited in the Sage Bionetworks NF Data Portal platform (https://www.synapse.org) with identification number syn21984813.

## Acknowledgements

This work was funded by grants from the NF Research Initiative (NFRI) and the St. Louis Men’s Group Against Cancer (to ACH). The authors would like to thank SpringWorks Therapeutics for supplying mirdametinib. Research reported in this publication was partially supported by the National Cancer Institute of the National Institutes of Health under Award Number U54CA224083. The content is solely the responsibility of the authors and does not necessarily represent the official views of the National Institutes of Health.

## Author Contributions

Himanshi Bhatia: Data curation; Formal Analysis; Investigation; Methodology; Validation; Visualization; Writing – original draft; Writing – review & editing

Alex T. Larsson: Data curation; Formal Analysis; Investigation; Methodology; Validation; Visualization; Writing – original draft; Writing – review & editing

Ana Calizo: Data curation; Formal Analysis; Investigation; Methodology; Validation; Visualization; Writing – original draft; Writing – review & editing

Kai Pollard: Data curation; Project administration; Methodology

Xiaochun Zhang: Data curation; Project administration

Eric Conniff: Data curation; Investigation; Methodology; Validation; Visualization; Writing - review & editing

Justin F. Tibbitts: Data curation; Investigation; Methodology; Validation; Writing – review & editing

Sara H. Osum: Methodology; Writing – review & editing

Kyle B. Williams: Data curation; Investigation; Methodology; Validation; Writing – review & editing

Ali L. Crampton: Data curation; Investigation; Methodology; Visualization; Writing – review & editing

Tyler Jubenville: Data curation; Formal analysis; Writing – review & editing

Daniel Schefer: Data curation

Kuangying Yang: Data curation; Visualization

Yang Lyu: Data curation; Formal Analysis; Investigation; Methodology; Validation; Visualization

Jessica Bade: Data curation

James C Pino: Formal analysis; Visualization

Sara J.C. Gosline: Conceptualization; Data curation; Formal analysis; Methodology; Writing – original draft; Writing – review & editing

Christine A. Pratilas: Conceptualization; Funding acquisition; Investigation; Project administration; Resources; Supervision; Writing – original draft; Writing – review & editing

David A. Largaespada: Conceptualization; Funding acquisition; Investigation; Project administration; Resources; Supervision; Writing – original draft; Writing – review & editing

David K. Wood: Conceptualization; Funding acquisition; Investigation; Project administration; Resources; Supervision; Writing – original draft; Writing – review & editing

Angela C. Hirbe: Conceptualization; Funding acquisition; Investigation; Project administration; Resources; Software; Supervision; Writing – original draft; Writing – review & editing

## The Paper Explained

### Problem

Multiple DNA anomalies are a unique characteristic of MPNST that make them difficult to treat in the clinic. An easy-to-use model system is needed that replicates the heterogenous nature of parental tumors and can be used to test candidate drugs.

### Results

We successfully established 13 PDX-MPNST lines and identified common somatic variants and chromosomal aneuploidy events. The PDX lines were further processed for assembly into 3D microtissues. Microtissues were categorized as “robust”, “good”, or “unusable” on the basis of viable cell count. Treatment of “robust” and “good” microtissues with commonly used MPNST therapy drugs (trabectedin, mirdametinib, and olaparib) identified mirdametinib as an effective single agent. We also observed synergy upon combination drug therapy. Treatment of corresponding PDX lines with the same drugs showed similar response pattern as the 3D microtissues.

### Impact

We report the development of a novel 3D microtissue platform that replicates the drug response pattern of *in vivo* PDX models. The data suggest that our model will be beneficial for future drug screening studies.

## Expanded View Figure legends

**Figure EV1. Dissociated PDX tumor cells rarely form spheroids**

A. Representative brightfield images of dissociated PDX tumor cells at two days of growth either beginning to aggregate or remain dispersed in collagen-free environment. Scale bars: 200μm.

B. Representative brightfield and fluorescent images showing either spheroid formation (MN-2) or continued cell aggregation (JH-2-031) after 7 days in culture in collagen-free environment. Cells were stained with calcein AM (live) and DRAQ5 (total). Scale bars: 200μm.

**Figure EV2. Some MPNST PDX cells fail to grow as 3D microtissues**

Representative brightfield and fluorescent images of several PDX 3D microtissues stained with calcein AM (live) and DRAQ5 (total) after two days in culture. Scale bars: 200μm.

**Figure EV3. Some PDX 3D microtissues remodel their collagen matrix**

Representative brightfield and fluorescent images showing MN-2 and JH-2-079-c microtissues remodeling the collagen matrix at different time points with the WU-225 matrix unchanged. Cells were stained with calcein AM (live) and DRAQ5 (total). Scale bars: 200μm.

**Figure EV4. PDX 3D microtissue data in response to drugs**

A. Dose response curves of four PDX 3D microtissues exposed to trabectedin combinations with either mirdametinib or olaparib at drug ratios of 1:2000 after two days in culture. Data points and error bars represent mean±SD, n≥10.

B. Area under the curve (AUC) of single agents and trabectedin combinations from panel A. Error bars drawn from bootstrap across the various experiments.

**Figure EV5. Differentially expressed genes across PDX**

Heatmap depicting 525 differentially expressed genes across 9 PDX engineered into microtissues categorized as “robust”, “good”, or “unusable”.

**Table EV1. List of differentially expressed genes based on microtissue quality**

