## Supplementary figures for "*Ex vivo* to *in vivo* model of malignant peripheral nerve sheath tumors for precision oncology"

### Slide 1
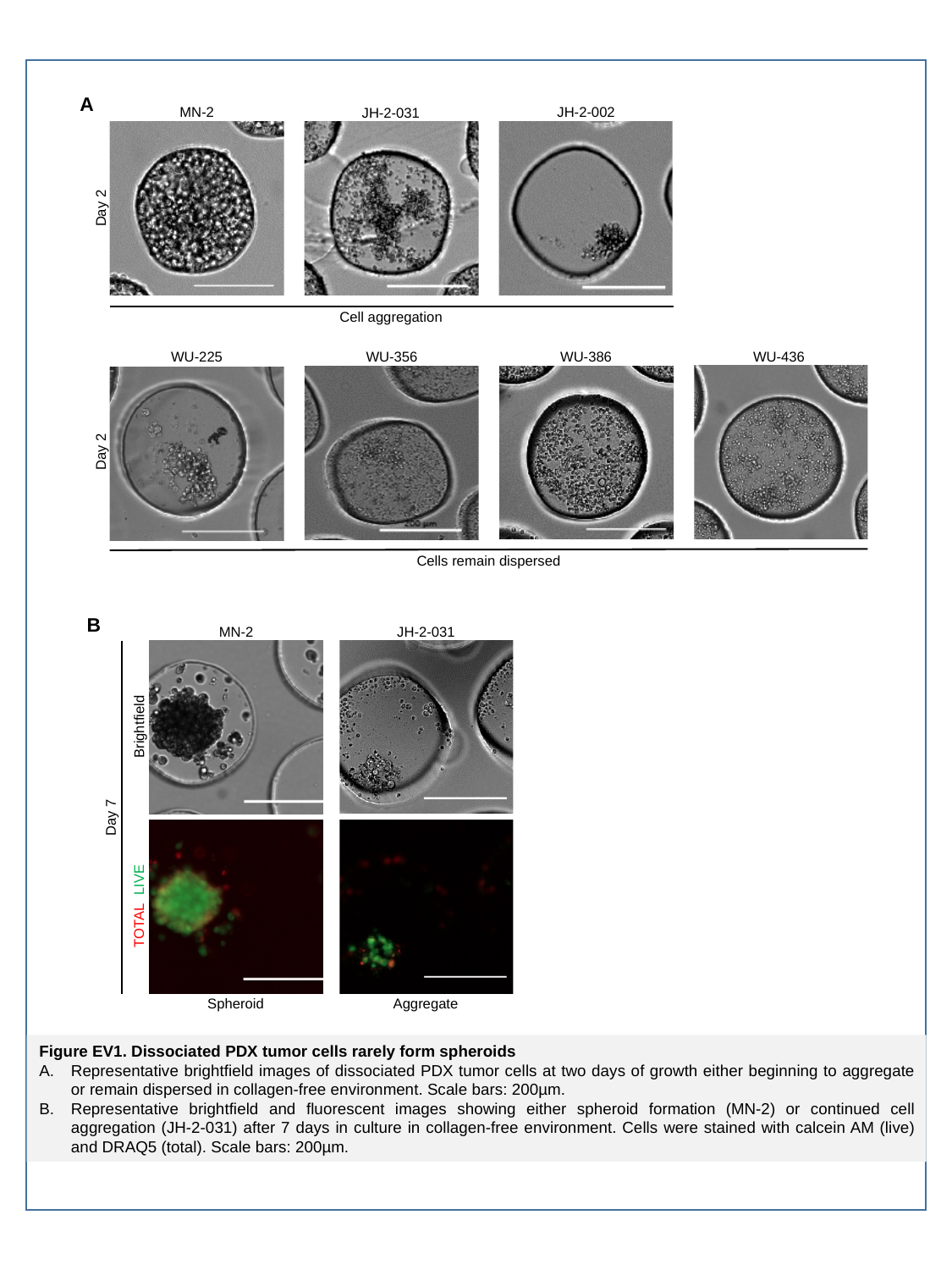

A
JH-2-002
MN-2
JH-2-031
Day 2
Cell aggregation
WU-356
WU-386
WU-436
WU-225
Day 2
Cells remain dispersed
B
JH-2-031
MN-2
Day 7
Brightfield
TOTAL LIVE
Aggregate
Spheroid
Figure EV1. Dissociated PDX tumor cells rarely form spheroids
Representative brightfield images of dissociated PDX tumor cells at two days of growth either beginning to aggregate or remain dispersed in collagen-free environment. Scale bars: 200µm.
Representative brightfield and fluorescent images showing either spheroid formation (MN-2) or continued cell aggregation (JH-2-031) after 7 days in culture in collagen-free environment. Cells were stained with calcein AM (live) and DRAQ5 (total). Scale bars: 200µm.

### Slide 2
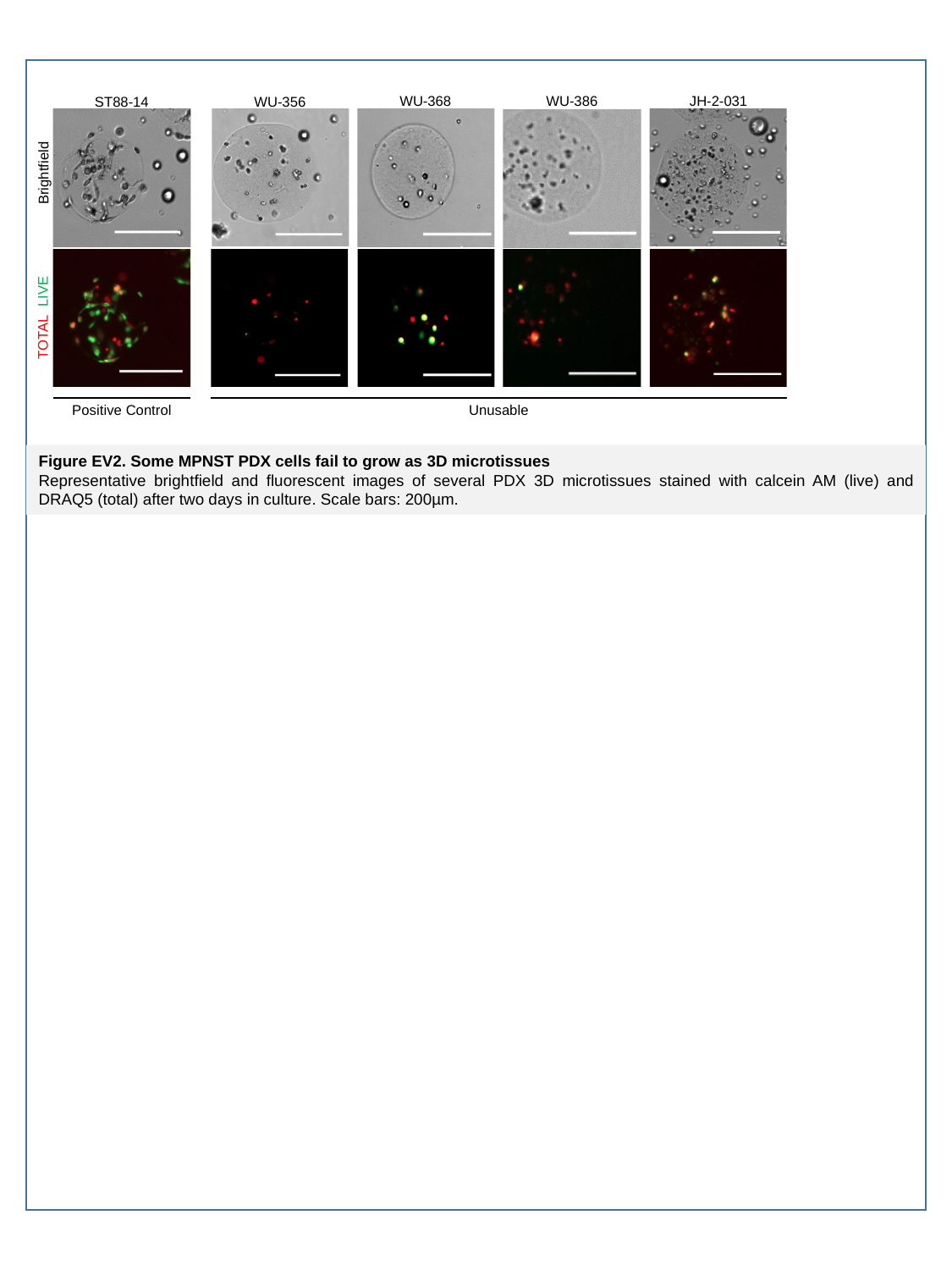

JH-2-031
WU-368
WU-386
WU-356
ST88-14
Brightfield
TOTAL LIVE
Positive Control
Unusable
Figure EV2. Some MPNST PDX cells fail to grow as 3D microtissues
Representative brightfield and fluorescent images of several PDX 3D microtissues stained with calcein AM (live) and DRAQ5 (total) after two days in culture. Scale bars: 200µm.

### Slide 3
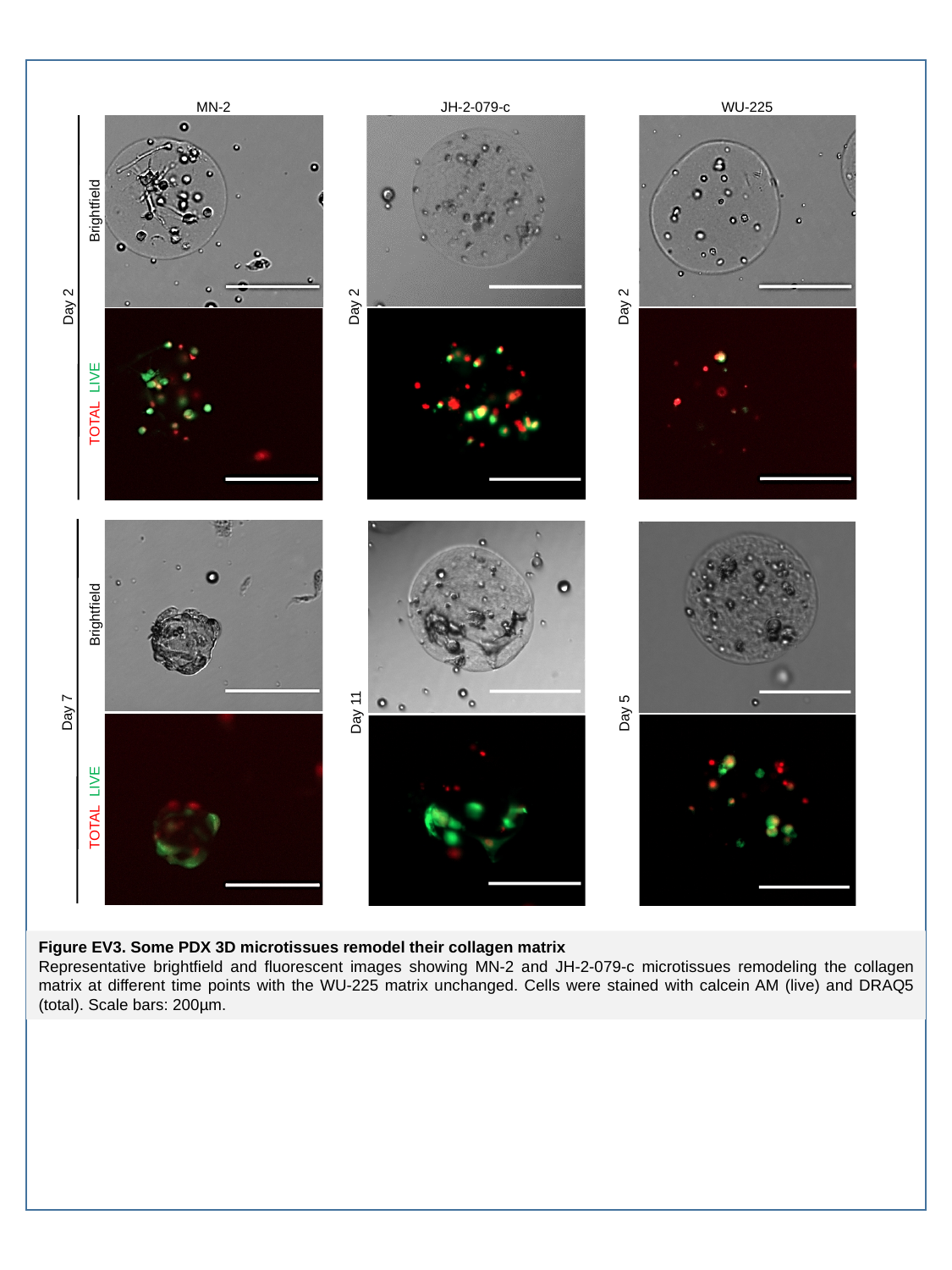

MN-2
Day 2
Day 7
Brightfield
TOTAL LIVE
Brightfield
TOTAL LIVE
JH-2-079-c
Day 2
Day 11
WU-225
Day 2
Day 5
Figure EV3. Some PDX 3D microtissues remodel their collagen matrix
Representative brightfield and fluorescent images showing MN-2 and JH-2-079-c microtissues remodeling the collagen matrix at different time points with the WU-225 matrix unchanged. Cells were stained with calcein AM (live) and DRAQ5 (total). Scale bars: 200µm.

### Slide 4
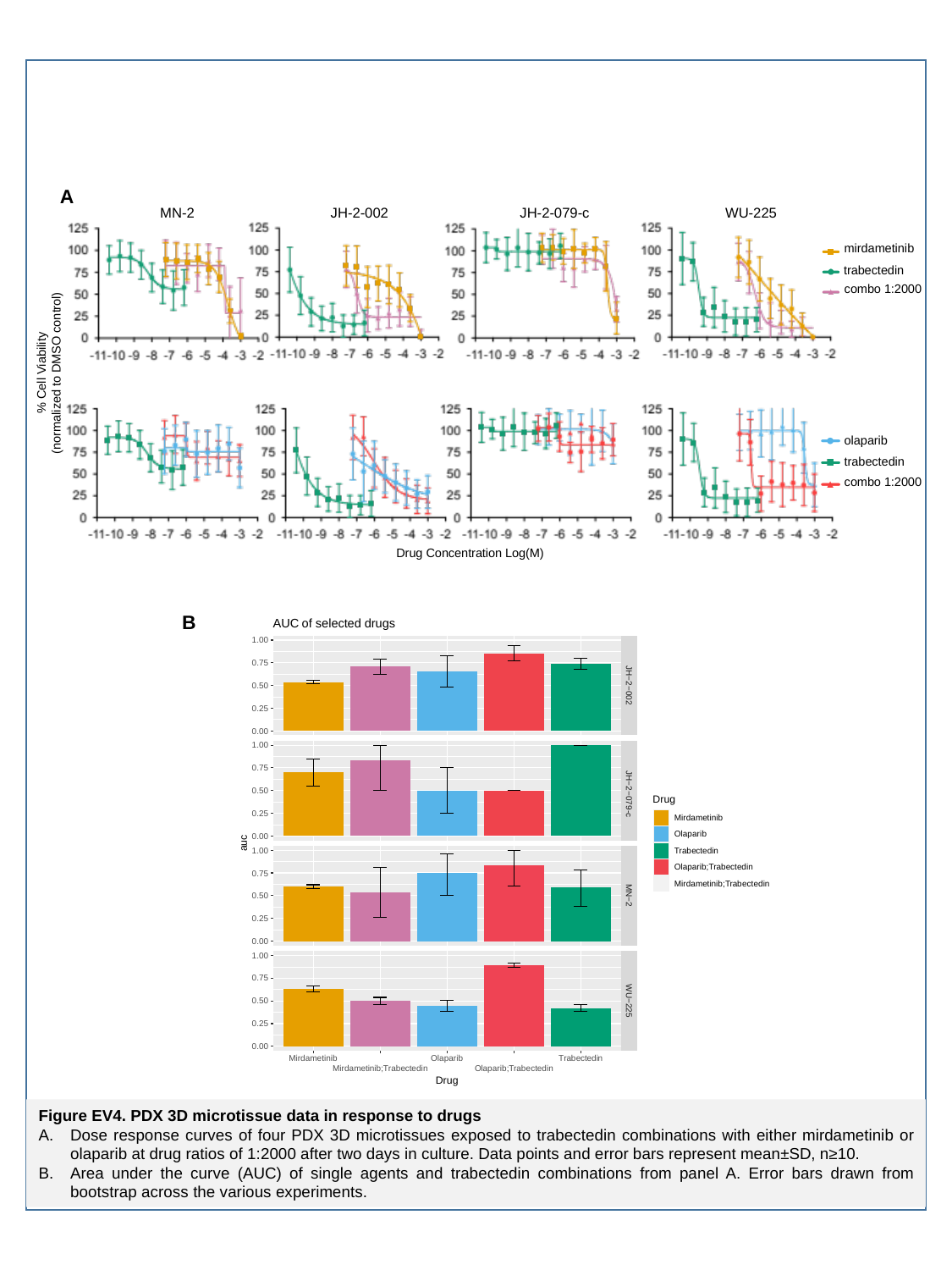

A
MN-2
JH-2-002
JH-2-079-c
WU-225
mirdametinib
trabectedin
combo 1:2000
% Cell Viability
(normalized to DMSO control)
olaparib
trabectedin
combo 1:2000
Drug Concentration Log(M)
B
Figure EV4. PDX 3D microtissue data in response to drugs
Dose response curves of four PDX 3D microtissues exposed to trabectedin combinations with either mirdametinib or olaparib at drug ratios of 1:2000 after two days in culture. Data points and error bars represent mean±SD, n≥10.
Area under the curve (AUC) of single agents and trabectedin combinations from panel A. Error bars drawn from bootstrap across the various experiments.

### Slide 5
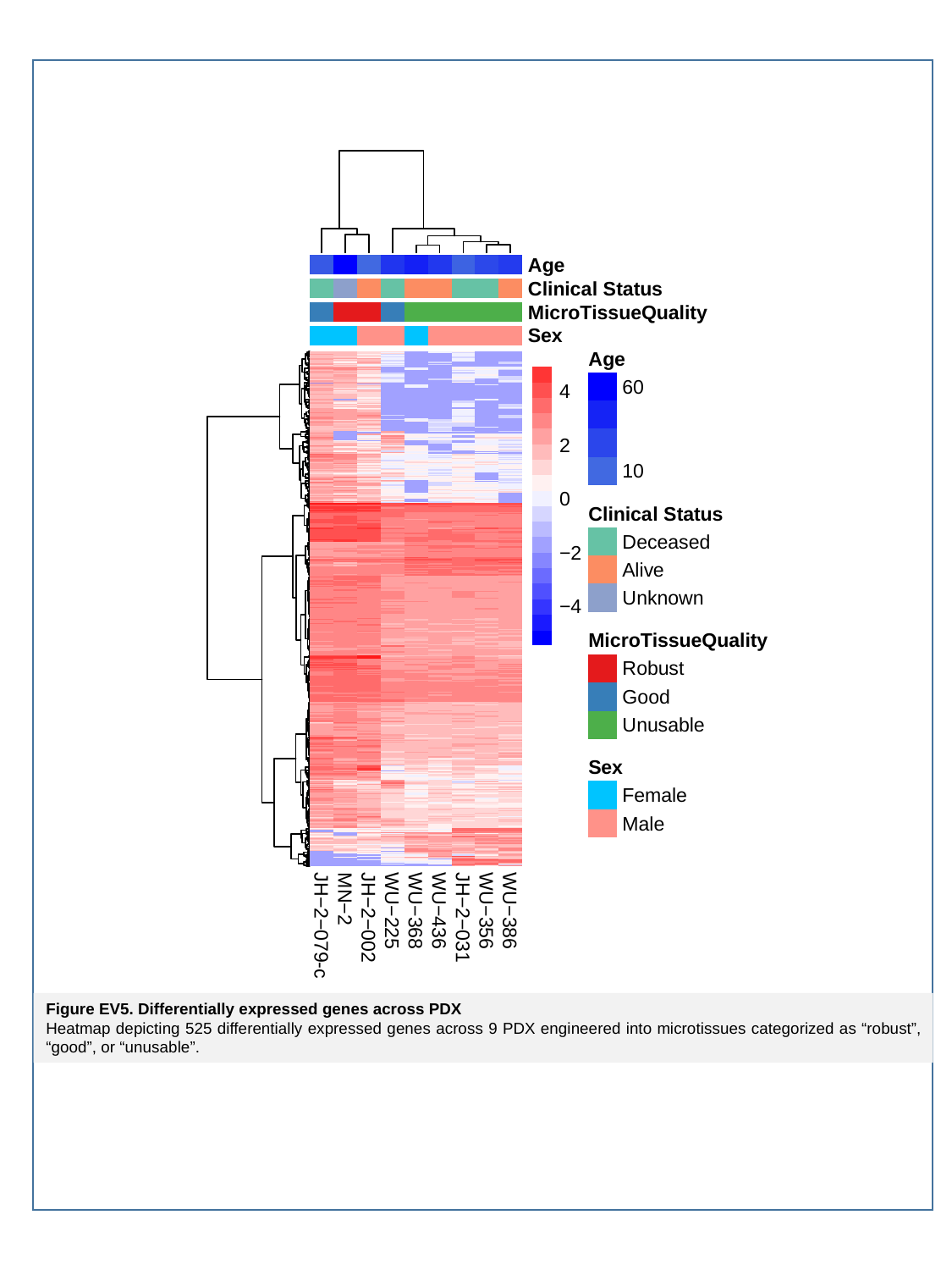

Figure EV5. Differentially expressed genes across PDX
Heatmap depicting 525 differentially expressed genes across 9 PDX engineered into microtissues categorized as “robust”, “good”, or “unusable”.
